# Exact sequence variants should replace operational taxonomic units in marker gene data analysis

**DOI:** 10.1101/113597

**Authors:** Benjamin J Callahan, Paul J McMurdie, Susan P Holmes

## Abstract

Recent advances have made it possible to analyze high-throughput marker-gene sequencing data without resorting to the customary construction of molecular operational taxonomic units (OTUs): clusters of sequencing reads that differ by less than a fixed dissimilarity threshold. New methods control errors sufficiently that sequence variants (SVs) can be resolved exactly, down to the level of single-nucleotide differences over the sequenced gene region. The benefits of finer taxonomic resolution are immediately apparent, and arguments for SV methods have focused on their improved resolution. Less obvious, but we believe more important, are the broad benefits deriving from the status of SVs as *consistent labels* with *intrinsic biological meaning* identified *independently from a reference database*. Here we discuss how those features grant SVs the combined advantages of closed-reference OTUs — including computational costs that scale linearly with study size, simple merging between independently processed datasets, and forward prediction — and of de novo OTUs — including accurate diversity measurement and applicability to communities lacking deep coverage in reference databases. We argue that the improvements in reusability, reproducibility and comprehensiveness are sufficiently great that SVs should replace OTUs as the standard unit of marker gene analysis and reporting.

## Introduction

High throughput sequencing of PCR-amplified marker genes has grown explosively over the past decade, especially as a means of taxonomically profiling microbial communities. Increasing use of marker-gene sequencing has been accompanied by increasing dataset sizes; this year, we can expect thousands of marker-gene studies to generate millions to billions of sequencing reads each.

The analysis of marker gene data customarily begins with the construction of molecular operational taxonomic units (OTUs): clusters of reads that differ by less than a fixed sequence dissimilarity threshold, most commonly 3% (Westcott & Schloss, 2015; Kopylova et al, 2016). The sample-by-OTU feature table serves as the basis for further analysis, with the observation of an OTU often treated as akin to the observation of a “species” in the taxonomic profiling application. Many methods for defining molecular OTUs have been proposed, but the most substantive distinction is between closed-reference methods — in which reads sufficiently similar to a sequence in a reference database are recruited into a corresponding OTU — and de novo methods — in which reads are clustered into OTUs as a function of their pairwise sequence similarities.

Recently, new methods have been developed that resolve sequence variants (SVs) from Illumina-scale amplicon data without imposing the arbitrary dissimilarity thresholds that define molecular OTUs (Eren et al, 2013; Tikhonov et al, 2015; Eren et al, 2015; Callahan et al, 2016a; Callahan et al, 2016b; Edgar, 2016). SV methods infer the biological sequences in the sample prior to the introduction of amplification and sequencing errors, and distinguish sequence variants differing by as little as one nucleotide. SV methods have demonstrated sensitivity and specificity as good or better than OTU methods, and better discriminate patterns of community similarity (Eren et al, 2013; Eren et al, 2015; Callahan et al, 2016a). The higher taxonomic resolution afforded by SV methods has self-evident benefits — for example, it is clearly useful to distinguish *Neisseria gonorrhoeae* from the many commensal *Neisseria* species in the human microbiota — and initial evaluation of SV methods has focused on their improved resolution. However, we argue here that the more important, and overlooked, advantage of SVs is that they combine the benefits for subsequent analysis of closed-reference and de novo OTUs: SVs are re-usable across studies, reproducible in future datasets, and are not limited by incomplete reference databases.

## Description of Methods

De novo OTU are constructed by clustering together sequencing reads that are sufficiently similar to one another. Many methods for constructing these clusters have been developed, but in all cases de novo OTUs are emergent features of a dataset, with boundaries and membership that depend on the dataset in which they are defined. This dataset dependence is not just a practical concern: the delineation of de novo OTUs depends on the relative abundances of the sampled community even in the limit of infinite sequencing depth and zero errors. As a result, de novo OTUs defined in two different datasets cannot be compared.

Closed-reference OTUs are properties of a reference database; each reference sequence in the database defines and labels an associated closed-reference OTU. Sequencing reads are assigned to closed-reference OTUs if they are sufficiently similar to the associated reference sequences. If the same reference database is used, closed-reference OTU assignments from independently processed datasets can be validly compared, a property we refer to as *consistent labeling*. However, biological variation that is not represented in the reference database is necessarily lost during assignment to closed-reference OTUs.

SVs are inferred by a de novo process in which biological sequences are discriminated from errors on the basis of, in part, the number of repeated observations of distinct sequences. As a result, SV inference cannot be performed independently on each read — the smallest unit of data from which SVs can be inferred is a sample. However, unlike de novo OTUs, *SVs are consistent labels* because SVs represent a biological reality that exists outside of the data being analyzed: the DNA sequence of the assayed organism. Thus, SVs that are inferred independently from different samples can be validly compared.

Figure 1 schematically represents the validity of de novo OTUs, closed-reference OTUs and SVs that were assigned from a common focal dataset. The x-axis represents all biological variation that exists at that genetic locus. The y-axis represents all amplicon data generated from that locus, and all future data that may be generated. The region of validity for each feature type is shaded.

**Figure 1:**
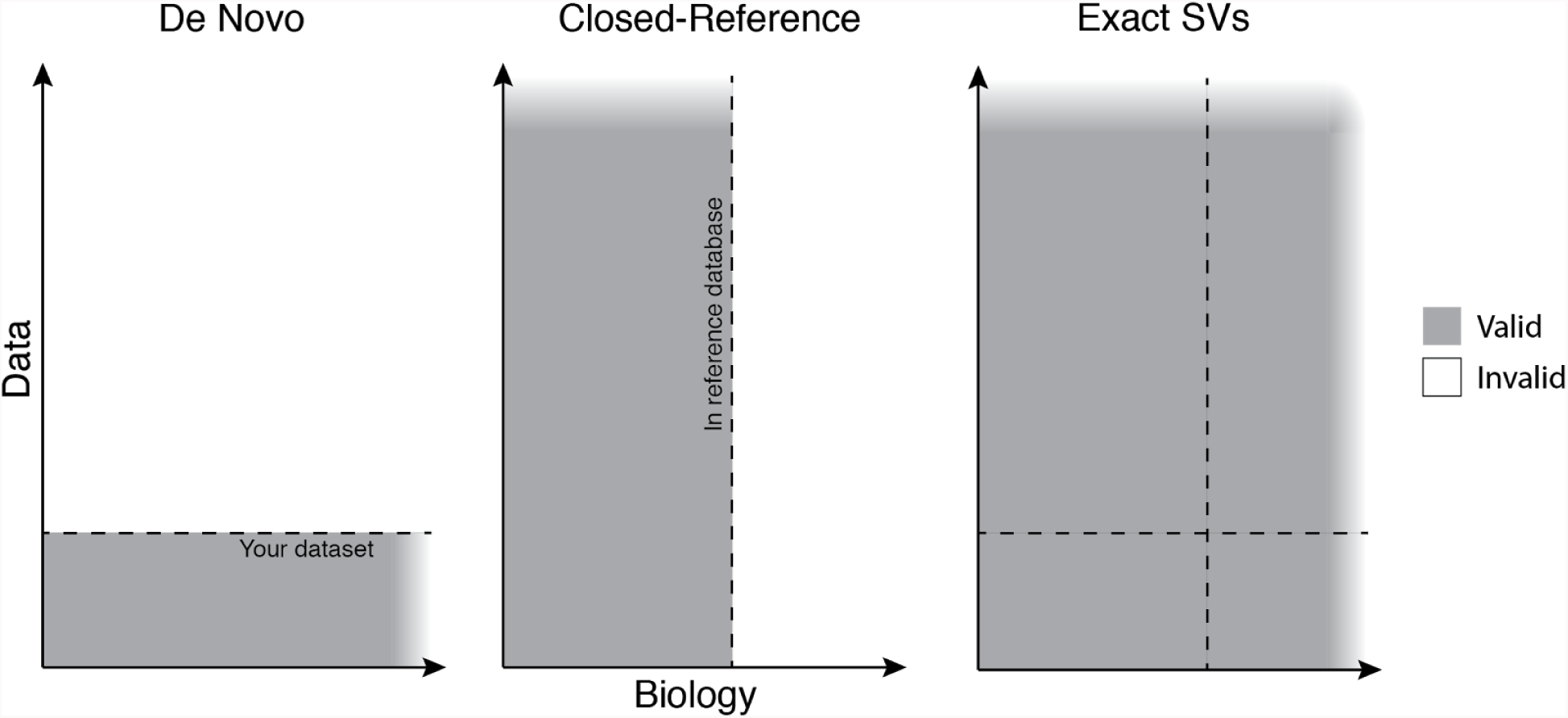
The extent of the validity of de novo OTUs, closed-reference OTUs and exact sequence variants (SVs) determined from a focal dataset.

This schematic emphasizes the limitations inherent to both classes of OTUs. De novo OTUs are invalid outside of the dataset in which they were defined. Closed-reference OTUs cannot capture biological variation outside of the reference database used in their construction. SVs transcend those limitations: SVs capture all biological variation present in the data, and SVs inferred from a given datasets can be reproduced in future datasets and validly compared between datasets.

## Practical consequences of consistent labels

### Computational tractability

Consistent labels allow the assignment of closed-reference OTUs to be split into the independent assignment of subsets of the data that are then merged together. De novo OTUs lack consistent labels so all data must be pooled for assignment, causing computational costs to scale quadratically with total sequencing effort and rendering common de novo methods prohibitively costly as study sizes increase (Rideout et al, 2014; Mahe et al, 2015).

SV inference cannot be performed on each read independently, but can be performed on each sample independently. As long as the sequencing effort devoted to individual samples remains tractable, independent inference by sample enables trivial parallelization and linear scaling of computation time with sample number, allowing SVs to be inferred from arbitrarily large datasets.

### Meta-analysis and replication

The growing number of marker-gene studies in similar environments creates opportunities for new analyses that combine studies for more power and generality. The consistent labeling of closed-reference OTUs and SVs allows per-study tables to be merged into a cross-study table. Meta-analysis is much more difficult with de novo OTUs, as the raw sequence data from each study must be compiled, pooled and reprocessed into new cross-study OTUs.

The absence of consistent labels makes replicating or falsifying previous results problematic, even impossible. Consider a significant association reported between the OTU “denovo123” and a condition-of-interest. This association cannot be tested in a new dataset, because “denovo123” does not exist in new datasets. Therefore, results from the analysis of de novo OTUs are only testable indirectly, by mapping OTUs onto consistent labels such as taxonomy or by reducing community composition to summary measures such as a diversity metric and then testing the results at that cruder level.

### Forward prediction

A major area of translational research is the use of microbial community composition as a predictive biomarker (Digiulio et al, 2015; Baxter et al, 2016). For example, the relative abundances of a set of OTUs or SVs might be used to predict a health condition. Predictive biomarkers can be constructed through statistical methods such as regression or by various machine learning methods, and their accuracy evaluated within the study by splitting the data into training and validation subsets (Callahan et al, 2016b). However, predictive biomarkers are only useful if they can be applied to new data. De novo OTUs exist only in the dataset in which the the predictor was trained and evaluated, so predictive biomarkers based on de novo OTUs can't predict from new data. SVs and closed-reference OTUs are applicable to new data, but closed-reference OTUs suffer from omitting predictive features absent from the reference database and thereby limiting predictive performance.

## Practical consequences of reference independence

### Diversity measurement

Reference databases are incomplete, so the assignment of reads to closed-reference OTUs removes that portion of the data that is unrepresented in the reference. This limitation is especially problematic if community diversity is of interest. The absence of unrepresented members of the community in closed-reference OTU tables can systematically skew diversity measures, potentially in a condition-dependent manner if some conditions are associated with a higher proportion of unrepresented taxa.

### Application across environments and genetic loci

The extent to which microbes inhabiting different environments are represented in reference databases varies greatly. In the best studied environments and genetic loci, such as 16S rRNA gene sequencing of the human gut, it is reasonable to expect upwards of 90% of sequencing reads to be assignable against a reference database. However, far fewer sequencing reads might be assignable in less well characterized environments or for genetic loci without built-up reference databases. SVs and de novo OTUs provide a much more accurate representation of the biological variation than closed-reference OTUs in less characterized environments or at less-studied loci.

### Guaranteed observation

The observation of a closed-reference OTU in a sample indicates that at least one sequencing read was sufficiently similar to the associated reference sequence to be mapped to its OTU. However, it does not guarantee that the reference sequence itself was observed — to the contrary, that is often not the case. Nevertheless, the often-absent reference sequence serves as the representative of the closed-reference OTU. This can be misleading when downstream analyses use the unobserved representative sequences to index into other sources of data.

### Changing references

Closed-reference OTUs are independent of the data, but they are wholly dependent on the set of reference sequences used, and even the order of those reference sequences (Westcott & Schloss, 2015). Therefore, closed-reference OTUs assigned against different reference databases are not comparable. In order to maintain comparability over time, reference databases must remain static, or closed-reference OTU tables must be regenerated whenever a new or upgraded reference database is adopted. Because SVs are consistent labels derived from the data, which does not change with time, they remain consistent into the indefinite future.

## Discussion

Molecular OTUs have served two different and orthogonal purposes. The first purpose is to translate taxonomic concepts developed in other systems into the context of high-throughput marker-gene sequencing of microbial communities by the ostensible equation of OTUs defined at certain thresholds with particular taxonomic levels (eg. 3% ribosomal OTUs are “like species”). The second purpose is to reduce the impact of amplicon sequencing error on measures of diversity and community composition by grouping errors together with the error-free sequence (Eren et al, 2016). A consequence of this often unacknowledged dual mandate is that OTUs struggle to serve both purposes well; the connection between OTUs and species is largely unfounded (Stackebrandt & Ebers, 2006), and the most common methods often output numbers of OTUs an order-of-magnitude higher than the number of strains present in mock communities (Kopylova et al, 2016).

There are ways to ameliorate some of the shortcomings of molecular OTUs. Open-reference OTU methods combine closed-reference OTU assignment with subsequent construction of de novo OTUs from the unassigned sequencing reads, gleaning the benefits of closed-reference OTUs without entirely sacrificing the unassignable portion of the data (Rideout et al, 2014). A clever computational approach has been developed that linearizes the computational time of a special case of de novo OTU assignment (single-linkage clustering with a linkage threshold of 1) allowing computational tractability on extremely large datasets (Mahe et al, 2015). Aggressive filtering and complete overlap between paired-end reads can reduce the rate at which OTU methods misinterpret sequencing artefacts as biological variation (Bokulich et al, 2013; Kozich et al, 2013; Edgar & Flyvbjerg, 2014).

Furthermore, the inference of exact SVs does not solve all problems. SV tables can only be merged if they cover the same gene region, so 16S data generated from different variable regions cannot be combined. The same genome can contain multiple SVs if there are multiple copies of the targeted genetic locus. Appealing terminology such as “resolution of exact sequence variants” does not eliminate the limitations inherent to representing a complex biological organism by a short genetic barcode. For example, while necessarily better than the customary 3% ribosomal OTUs, there is still no guarantee of ecological coherence or even monophyly among genomes with the same ribosomal SV (Berry et al, 2017).

Those caveats stated, the breadth of issues that SVs cleanly solve, and the more powerful and reproducible analyses that SVs enable, makes a dispositive case in our opinion for replacing OTUs with SVs. SVs have an intrinsic biological meaning, and correspond as closely as possible to the fundamental unit of microbial communities: the strain. SV inference from large marker gene datasets is both tractable and comprehensive. SVs improve the return-on-investment of marker-gene sequencing by better leveraging the corpus of such datasets for further discovery, especially in communities investigated in many studies like those inhabiting the human body. And the SV methods that are now available provide better resolution and accuracy than OTU methods (Eren et al, 2015; Callahan et al, 2016a).

For analysis to be reproducible the fundamental units must be reproducible, and de novo OTUs are not. For analysis to be comprehensive the fundamental units must be comprehensive, and closed-reference OTUs are not. Replacing OTUs with SVs makes marker gene sequencing more precise, reusable, reproducible, and comprehensive. We believe that SVs should be the standard way that marker gene data is processed and reported going forward.

